# ReadFilter - Filtering reads of interest for quicker downstream analysis

**DOI:** 10.1101/266080

**Authors:** Kim Lee Ng, Thor Bech Johannesen, Mark Østerlund, Kristoffer Kiil, Paal Skytt Andersen, Marc Stegger

## Abstract

Whole-genome sequencing is becoming the method of choice but provides redundant data for many tasks. ReadFilter (https://github.com/ssi-dk/serum_readfilter) is offered as a way to improve run time of these tasks by rapidly filtering reads against user-specified sequences in order to work with a small fraction of original reads while maintaining accuracy. This can noticeably reduce mapping time and substantially reduce *de novo* assembly time.

## Introduction

In the last decade, whole-genome sequencing (WGS) has changed from a high-profile research tool to become the gold standard of many reference laboratories around the world. Data generated from it is now used concurrently to understand gene flow, virulence potential, resistance problems, outbreaks and population dynamics, and spans between identifying ward-specific outbreaks to the unraveling of worldwide epidemic clones. Often these investigations include searches for specific genes or gene variants of interest (e.g. multilocus sequence typing (MLST) genes (Maiden et al., 1998), resistance genes, virulence genes, core genome MLST genes, etc.) to aid both researchers and clinicians in their interpretation of the sequence data. These genes or regions typically comprise only a fraction of the generated reads.

Here we present ReadFilter (https://github.com/ssi-dk/serum_readfilter), a Linux command-line tool which rapidly filters reads for downstream applications given any set of reference sequences to filter against. It produces an output of sequence reads that are potentially from the genes of interest and which constitute only a fraction of the input WGS data. This filtering of reads is important to achieve reduced run time of many applications used in research groups, core sequencing facilities and centers running publically available typing tools that have to handle large quantities of WGS data sets. Tools such as ARIBA (Hunt et al., 2017) also perform filtering with their analysis workflow. ReadFilter differentiates from ARIBA by separating it from the analysis workflow, and as the output are the same file type as the input it allows quick implementation into existing fastq-based workflows.

## Methods

ReadFilter works by creating a database converted from user input reference set (fasta sequences) of interest and taking in WGS paired-end reads. These reads are then checked against the database and returns an output containing the filtered set of read files which share a specified degree of similarity with the database. ReadFilter accomplishes this by utilizing either the Kraken (Wood & Salzberg, 2014) or Kaiju (Menzel, Ng, & Krogh, 2016) tools. Kraken filters reads that share a k-mer with those in the reference set and Kaiju filters reads which share an amino acid segment with the reference set. These tools were both designed for metagenomics classification of samples so applying their logic for filtering WGS samples simplifies their approach as the approach do not require classification but only determine if a sequence classifies or not. Depending on the sensitivity desired the k-value used to generate the kraken database, or the seed size for Kaiju, can be adjusted with a smaller k-value reducing precision while increasing sensitivity (or vice-versa). As the database is expected to be smaller than the genome, generation of the database is very rapid (seconds). Additionally, the amount of information in the filtered reads can be further reduced through subsampling or normalization (an included option to normalize the data using BBNorm (Bushnell, n.d.) is available. The inverse can also be achieved to filter out reads which have any potential to belonging to the database (a contamination/identifier filter).

ReadFilter was tested against available outbreak datasets (Page et al., 2017) using a Red Hat Enterprise Linux Server v7.4 cluster with each job given 1 CPU and 32 Gb of memory. The raw reads were run through srst2 v0.2.0 (Inouye et al., 2014) and SPAdes v3.11.0 (Bankevich et al., 2012) then mlst v2.9 (Seemann, n.d.) both with and without application of ReadFilter. Timings (Table 1) were also compared with the addition of BWA-MEM v0.7.17 (Li & Durbin, 2009).

**Table 1.** Timings of software against 85 samples (from four different species)

| Software | Filter time | Run time | Total time |
| --- | --- | --- | --- |
| mlst (filtered)<sup>1</sup> | 250 | 1.4 (+34.5) | 285.9 |
| mlst (unfiltered)<sup>1,2</sup> | - | 1.9 (+2873) | 2874.9 |
| srst2 (filtered) | 250 | 2147.7 | 2397.7 |
| srst2 (unfiltered)<sup>2</sup> | - | 2380.2 | 2380.2 |
| BWA-MEM (filtered) | 250 | 11.4 | 261.4 |
| BWA-MEM (unfiltered) | - | 326.1 | 326.1 |

All times are in minutes
<sup>1</sup>The time to assemble with SPAdes before running the application was 34.5 and 2873 minutes and included separately
<sup>2</sup>Timings taken from (Page et al., 2017)

## Results

Running ReadFilter reduced the read count of samples for MLST genes by 3-4 orders of magnitude (Figure 1).

**Figure 1.**
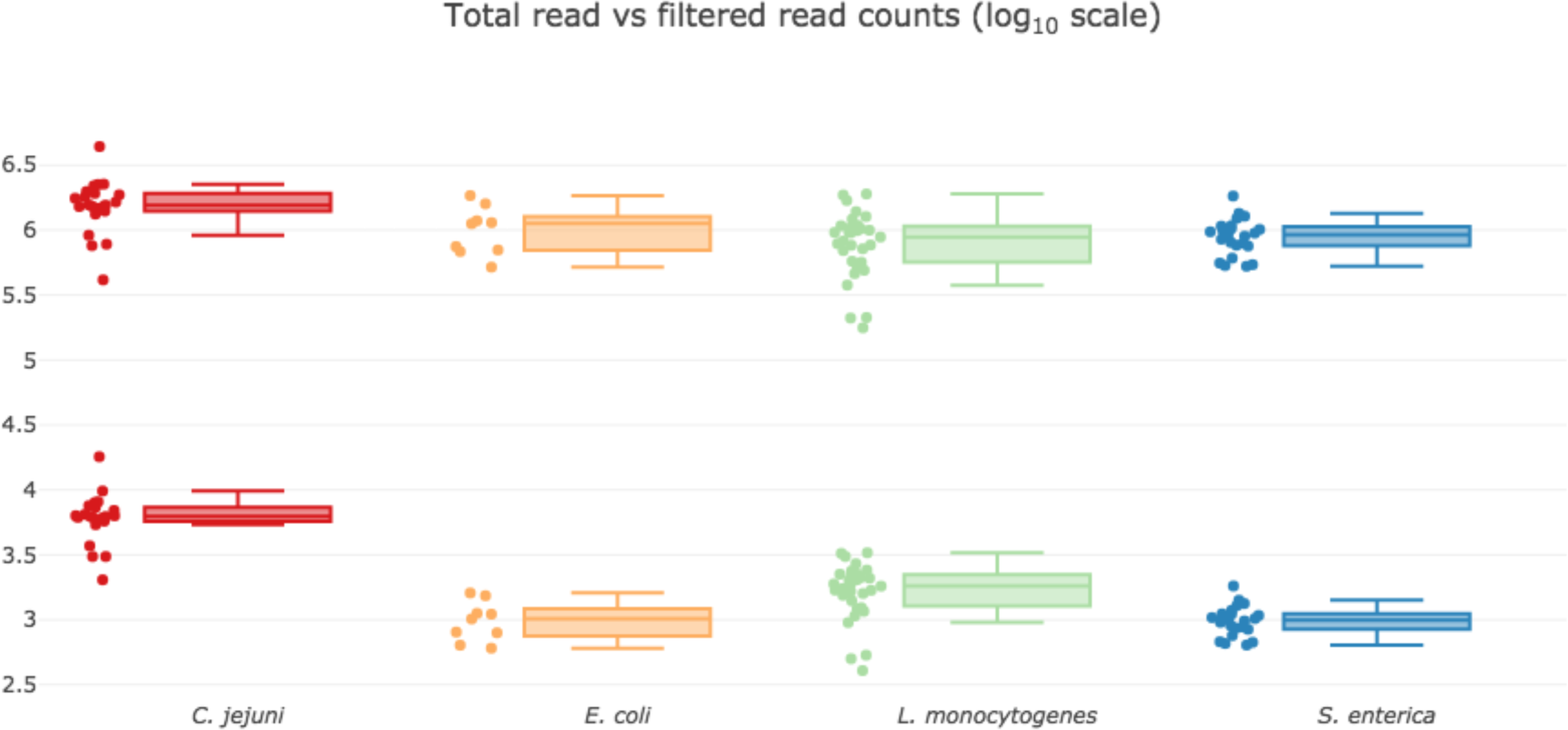
Reduction in reads using ReadFilter on four different species. Shows the reduction of reads each sample has within a clustered species when filtered against the MLST genes of interest with box plots. The upper box plots represent the number of paired raw read counts and the bottom box plots the filtered read counts obtained from ReadFilter as clustered on species.

Results for all 85 samples were in concordance when using the filtered reads for MLST at 100% using srst2 and matched 97.6% using SPAdes and then mlst. For the two discrepant samples no allelic calls were made as the genes of interest was split onto two contigs but otherwise matched 100%. MLST-based methods mlst (contigs) and srst2 (mapping) were selected as representatives for benchmarking time and BWA-MEM to compare an additional mapper. There was an order of magnitude speed increase when doing assembly with SPAdes on the reduced data set for running software mlst. While there was a slight reduction in speed for running srst2 the gains are lost with the time taken for the filtering step. This is likely due to how srst2 maps reads to all references so that any increase in speed with dealing with a lower read count it is fairly negligible by the time required to map to all potential candidates. In contrast, BWA-MEM had notable speed gains from the addition of the filtering step. While benchmarking was done to match resource use in existing benchmarks (Page et al., 2017), the filter step parallelizes efficiently with multiple processors while downstream programs may not meaning practical time spent filtering is likely to be a smaller portion of runtime.

## Discussion

Searches for specific genes or gene variants is a routine process and an integral part of microbial whole-genome sequenced-based analyses. Such analyses may seem rapid for occasional users, however for a large number of users these tasks are cumulatively very time consuming. By filtering potential reads out of the entire generated sequence dataset, a substantial reduction in computation time can be achieved. This approach can similarly be applied in other areas such as examining specific gene/exon of interest in RNAseq, and searching for genes of interest in metagenomics samples. Depending on specific requirements filtering techniques such as an Amino Acid BWT with Kaiju and a k-mer-based Kraken are available. Other filters can be added if necessary depending on requirements though with short reads, Kraken should work for most foreseeable cases. In reference to other MLST-based approaches, methods which do not filter reads efficiently can benefit from ReadFilter. Based on our results this can be realized in speed gains on *de novo* assembly approaches.

Reducing the required reads, especially if the reads need to be used for multiple applications, can lead to significant speed gains and a reduction of computational resources while providing a similar end result. Institutions can benefit from pre-filtering of reads for downstream analysis with minimal impact on existing workflows due to the input being the same file type as the output. Existing workflows based around receiving all reads can also reduce data transfer if users pre-filter their data before submission. Approaches which may have been computationally too expensive on all reads may now be feasible on a reduced read set. For applications with larger databases such as cgMLST, which can comprisê50% of the genome (de Been et al., 2015), species depending, there is still a potential gain of removing the remaining ∼50% of the reads and perform more efficient assemblies. Alternatively, returning the reads which do not match the database provides a smaller working set for accessory genome analyses, as well as resistance or virulence marker identification. As the number of reads is often dramatically reduced using ReadFilter, other mapping approaches to provide additional information including heterozygous positions or examining flanking regions can be exploited, information that contig based approaches do not allow.

Additional filtering or classification information can be beneficial to certain tasks by filtering the filtered data set to obtain a subset of the data or classifying the filtered reads. This would apply to tasks such as multi-gene mapping where initial filtering would be done on all gene candidates, followed by additional filtering on individual gene clusters to reduce candidate reads for each cluster. For diverse gene candidate sets this would help reduce the read count for each individual gene or gene cluster, therefore improving mapping time. While the method and approach described here is based on Illumina reads, it contains no limitation to this platform and the approach can work on all short-read platforms as well as long read data such as from Oxford Nanopore’s MinION platform or PacBio reads. Due to the higher error rate currently observed in long reads, utilizing Kaiju may be a more appropriate filtering approach to use depending on how the errors are distributed in the reads.

## Conclusions

The gains from targeted sequence read filtering are currently underused and would immediately benefit many users of databases referring to a small fraction of the genome. Filtering through ReadFilter allows only relevant reads to match a sequence database of interest and can provide a reduction of up to four magnitudes of reads to be considered for downstream applications. As ReadFilter produces identical file types (fastq) of the provided input files, it can easily be deployed with minimal changes to existing pipelines and workflows.

## Data bibliography

1. Ng, K L. https://github.com/ssi-dk/serum_readfilter (2018)

2. Katsz, L. https://github.com/WGS-standards-and-analysis/datasets (2018)

